# Inhibition of amyloid formation of the Nucleoprotein of SARS-CoV-2

**DOI:** 10.1101/2021.03.05.434000

**Authors:** Einav Tayeb-Fligelman, Xinyi Cheng, Christen Tai, Jeannette T. Bowler, Sarah Griner, Michael R. Sawaya, Paul M. Seidler, Yi Xiao Jiang, Jiahui Lu, Gregory M. Rosenberg, Lukasz Salwinski, Romany Abskharon, Chih-Te Zee, Ke Hou, Yan Li, David R. Boyer, Kevin A. Murray, Genesis Falcon, Daniel H. Anderson, Duilio Cascio, Lorena Saelices, Robert Damoiseaux, Feng Guo, David S. Eisenberg

## Abstract

The SARS-CoV-2 Nucleoprotein (NCAP) functions in RNA packaging during viral replication and assembly. Computational analysis of its amino acid sequence reveals a central low-complexity domain (LCD) having sequence features akin to LCDs in other proteins known to function in liquid–liquid phase separation. Here we show that in the presence of viral RNA, NCAP, and also its LCD segment alone, form amyloid-like fibrils when undergoing liquid–liquid phase separation. Within the LCD we identified three 6-residue segments that drive amyloid fibril formation. We determined atomic structures for fibrils formed by each of the three identified segments. These structures informed our design of peptide inhibitors of NCAP fibril formation and liquid–liquid phase separation, suggesting a therapeutic route for Covid-19.

**One Sentence Summary:** Atomic structures of amyloid-driving peptide segments from SARS-CoV-2 Nucleoprotein inform the development of Covid-19 therapeutics.

## Introduction

The Nucleoprotein (NCAP) of Severe Acute Reparatory Syndrome Coronavirus 2 (SARS-CoV-2) functions in viral replication by packaging the genomic RNA (gRNA) and participating in virion assembly(*1–9*). In the presence of RNA, NCAP forms liquid–liquid phase separation (LLPS) which increases RNA local concentration in cells(*1–8*). This LLPS was postulated to be driven by the NCAP’s intrinsically disordered regions (IDRs); the longest of which is the 75-residue low-complexity domain (LCD) bridging between the protein’s RNA binding domain (RBD) and its dimerization domain (DD) (Figure 1A,B) (*4, 5, 7*). Similar LCDs in other RNA binding proteins (RBPs), including the human Fused in Sarcoma (FUS), TDP-43 and hnRNPA2, are known to drive LLPS, accompanied by amyloid formation and droplet maturation into solid-like hydrogels (*10, 11, 20, 21,12–19*). Amyloid fibrils are structured protein aggregates amenable to structure-guided inhibitor design(*22*). We therefore hypothesized that NCAP is capable of forming amyloid, and that manipulation of this process, via designed fibrillation modulators, may hinder the protein’s function, and thereby inhibit viral replication.

**Fig. 1.**
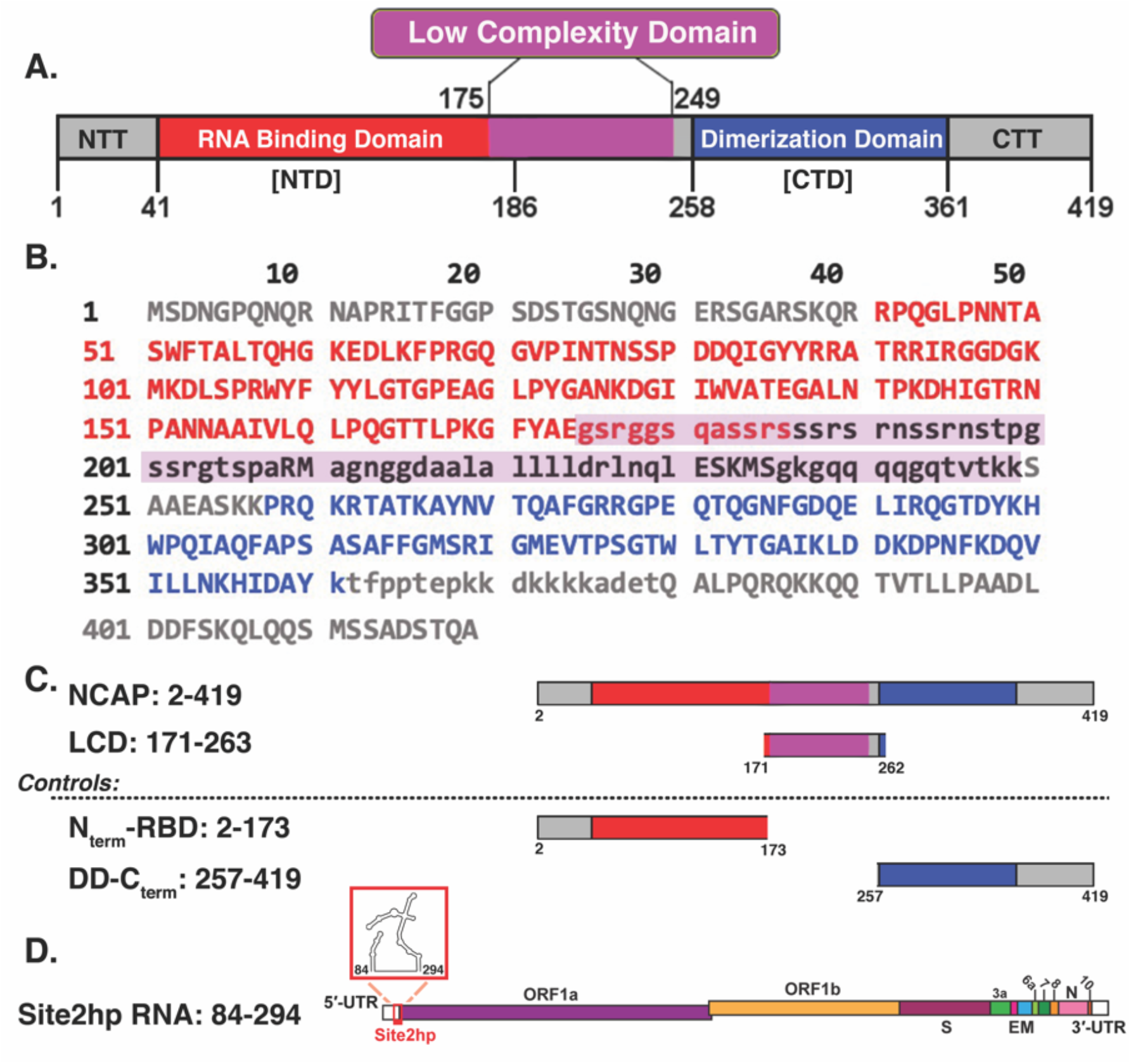
Overview of NCAP domain organization and sequence, and of protein and RNA constructs used in this study. (A) Domain designations of the SARS-CoV-2 NCAP: N-terminal tail (NTT in gray), RNA binding domain (RBD in red); a low complexity domain (LCD purple; residues 175-249), a dimerization domain (DD in blue); the C terminal tail (CTT in gray). (B) Amino acid sequence of the NCAP colored according to the domains presented in (A). Residues considered as low complexity are written in lower-case letters. (C) Protein constructs and controls used in this study, abbreviated NCAP, LCD, N_term_-RBD and DD-C_term_. The first N-terminal residue in all of those constructs (residue #1) is a threonine residing from cleavage of the 6xHis-SUMO tag during protein purification. (D) A schematic representation of SARS-CoV-2 genome and the location of hairpin site 2 (Site2hp) RNA used in this study.

## Results

Induced formation of NCAP LLPS by RNA depends on RNA length and concentration (*2, 7, 8*). To define the factors that govern the postulated NCAP fibril formation in the presence of RNA, we examined the protein-RNA interactions of NCAP, and its segments. The proteins chosen for this study are the full-length NCAP, its LCD segment, and two controls spanning the DD to C-terminus segment (DD-C_term_) and the N-terminus to RBD (N_term_-RBD) (Figure 1C). A recent study showed that the 5’- and 3’-ends of the SARS-CoV-2 viral genome promote the formation of NCAP LLPS, and two prominent NCAP-binding sites have been identified in the 5’ region(*7*). These sites are characterized as single-stranded RNA flanked by structured elements. In our study, we selected one of them, called Site2hp, that contains a 22-nucleotide A/U-rich unstructured strand sequence and its neighboring hairpins (Figure 1D). Hairpins were included for extended RNA length, and to allow potential interactions between the protein and the structured RNA regions. We assayed the mobility of the mixtures of Site2hp RNA with NCAP, or its individual segments, relative to the mobility of the RNA alone and protein alone using size exclusion chromatography (SEC) and gel shift assays. Site2hp demonstrated binding to NCAP in 2:1, and more so in 4:1, protein:RNA molar ratio, as evident from both assays (Figure 2 & S1). In the gel shift assay, DD-C_term_ exhibited partial shift of the RNA band only in the 4:1 protein:RNA molar ratio, indicative of DD-C_term_-Site2hp binding, but presumably of weaker affinity than those of the NCAP-Site2hp. No strong binding was detected with the N_term_-RBD or the LCD segments under those experimental conditions (Figure 2). These results overall indicate that Site2hp contains several NCAP binding sites, therefore requiring excess of NCAP for strong Site2hp binding, and that the different protein domains function in the full length NCAP in a cooperative mode to promote RNA-binding as was previously described for the NCAP of SARS-CoV(*23*).

**Fig. 2.**
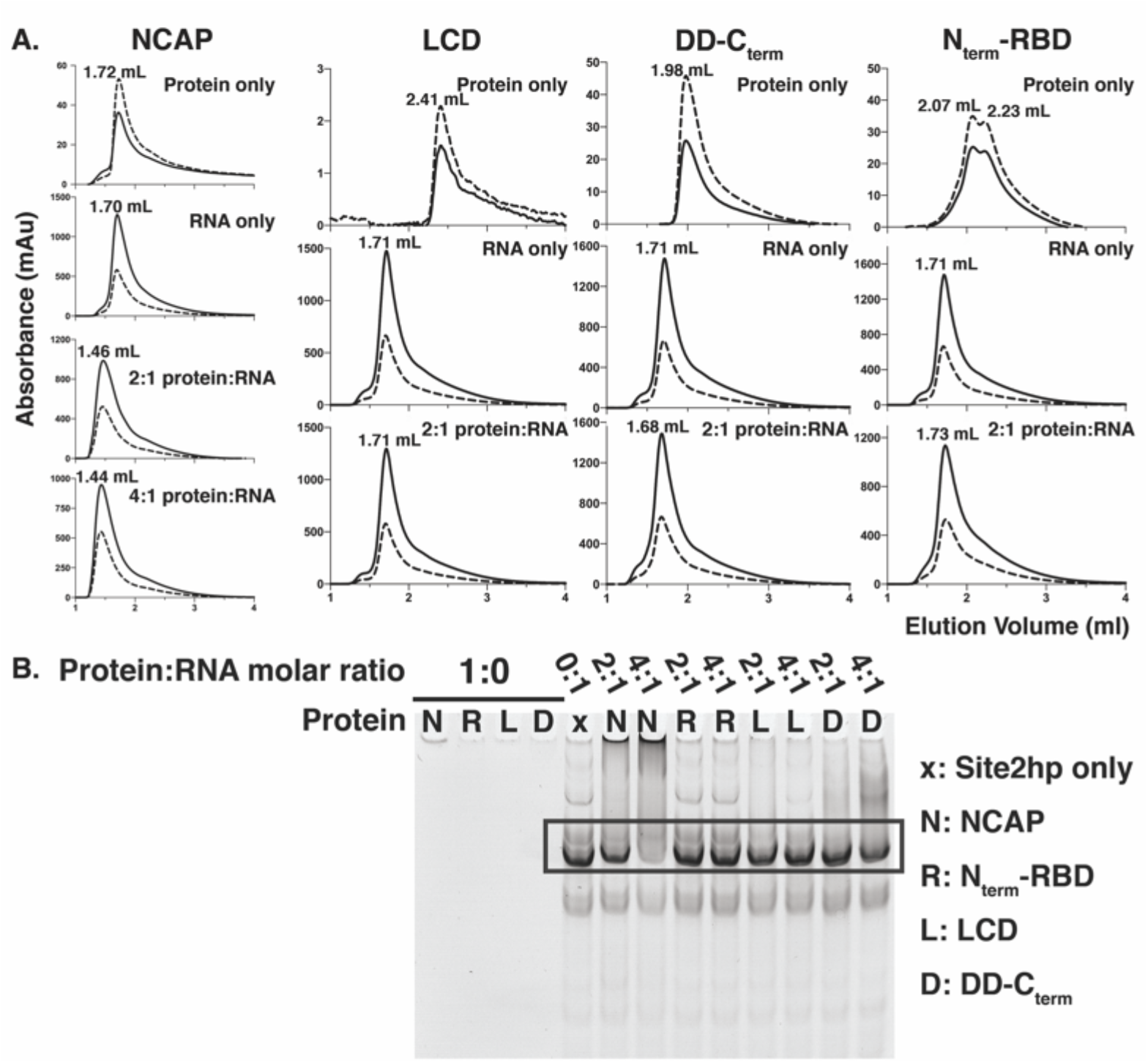
Hairpin Site2 (Site2hp) RNA and its binding to NCAP and its segments as detected by size exclusion chromatography (SEC) and native gel analysis. A) SEC chromatograms of the free NCAP protein or its segments, free Site2hp RNA, and corresponding mixtures. Dashed lines show absorbance at 280 nm; solid lines at 260 nm. Left: NCAP binds to Site2hp RNA in the tested 2:1 and 4:1 protein:RNA molar ratios. Free NCAP protein concentration is 24 μM, Site2hp concentration is 12 μM, and 2:1 and 4:1 protein:RNA mixtures contain 24 μM and 48 μM of the NCAP respectively. Middle left, Middle right & right: The LCD, DD-C_term_, and N_term_-RBD segments respectively exhibit no strong binding to Site2hp RNA. Segment’s concentration is 24 μM, Site2hp RNA concentration is 12μM. B) Native gel mobility shift assay detecting interactions between Site2hp RNA and the NCAP protein or its segments. The proteins NCAP (N), N_term_-RBD (R), LCD (L), DD-C_term_ (D) were separately mixed with Site2hp RNA in 1:0 (no RNA), 0:1 (x; Site2hp only), 2:1 and 4:1 protein:RNA molar ratios. Free intact Site2hp RNA is depicted within the black box. Shift of the free RNA band is indicative of protein-RNA binding. NCAP demonstrates some binding to Site2hp in 2:1 and 4:1 protein:RNA ratios, with the most pronounced binding shown for the 4:1 mixture. The NCAP-RNA complex is mostly trapped in the well region of the gel, typical of amyloid fibrils and other extremely large complexes. DD-C_term_ binding to the RNA is detectable only in the 4:1 protein:RNA mixture, whereas the LCD and N_term_-RBD does not demonstrate any strong binding at tested protein:RNA ratios. These results are overall in agreement with the protein-RNA binding detected by SEC for the corresponding tested mixtures shown in A.

Site2hp RNA promotes amyloid fibril formation of LCD and, to a lesser extent, of NCAP and DD-C_term_ as judged by fluorescence of Thioflavin T (ThT), an amyloid specific dye. Low complexity regions appear to have facilitated aggregation, since all three of these NCAP constructs contain them (75 residues in NCAP and LCD, and 19 residues in DD-C_term_; Figure 1B). In contrast, the N_term_-RBD segment contains no low complexity region and produced no ThT signal throughout the ~33h of measurement. RNA-deprived NCAP constructs also produced no ThT signal for ~33 h (Figure 3 & S2). These trends in aggregation propensity are echoed by a parallel set of experiments monitoring turbidity, which is free of ThT reporter, and indicative also of other forms of aggregation, including LLPS formation (Figure 3). Electron micrographs of LCD-Site2hp show abundant, elongated, unbranched fibrils typical of amyloid (Figure S3). Overall, these observations indicate that NCAP, its LCD, and to some extent the DD-C_term_ segment, are prone to form amyloid. Notably, amyloid formation is significantly stimulated, and morphologically altered (Figure S3) by protein-RNA interactions.

**Fig. 3.**
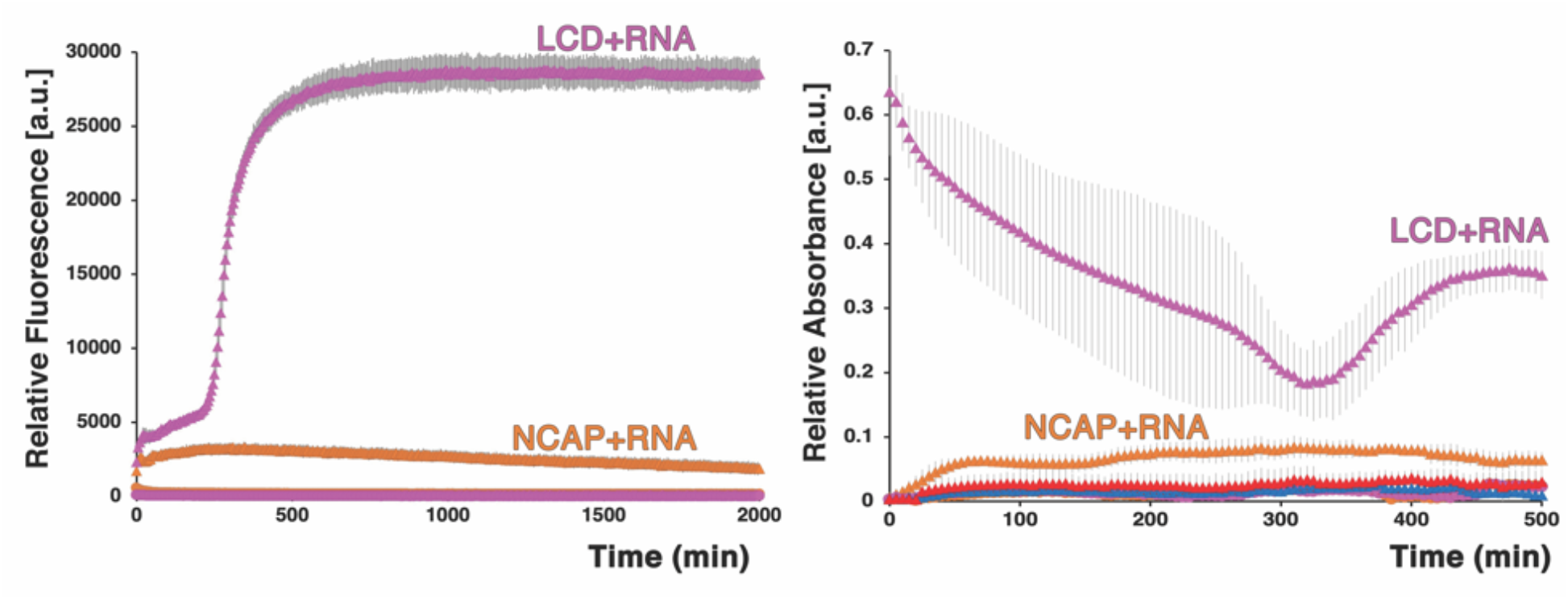
Site2hp RNA enhances the rate of amyloid-like fibril formation of NCAP its LCD segment, and to a lesser extent of its DD-C_term_ segment, as shown by Thioflavin-T fluorescence and by turbidity. Left: Relative fluorescence of protein-(buffer/RNA)-ThT mixtures compared to that of the protein deprived blank mixtures. Notice that the LCD is most strongly affected by RNA, followed by the NCAP. LCD alone (magenta circles), and NCAP alone (orange circles) are virtually at baseline. DD-C_term_ is only mildly affected by RNA, whereas N_term_-RBD showed no fibril formation (Figure S2). Right: Turbidity of solutions also shows that LCD and the NCAP proteins are most strongly affected by RNA addition. The relative absorbances for LCD, DD-C_term_, N_term_-RBD, and NCAP in the absence of RNA, as well as DD-C_term_ +RNA, and N_term_-RBD+ RNA, are essentially all at baseline. Both ThT and turbidity experiments were repeated three times in three different days. Representative curves are shown. Error bars represent standard deviation (SD; some too small to be visible).

Amyloid and LLPS formation are correlated in mixtures of NCAP, and its LCD, with appropriate Site2hp RNA concentrations, as visualized *in vitro* by fluorescence microscopy with the amyloid fluorescent dye Thioflavin-S (ThS). ThS positive LLPS droplets of ~2-5 μm in diameter were observed of the NCAP protein when mixed with Site2hp RNA in 40:1 protein:RNA molar ratio (0.75 μM RNA). Those droplets were also positive to propidium iodide (PI), a nucleic acid indicator, demonstrating that the protein fibrils and RNA are co-localized into those droplets (Figure 4A). A 4:1 NCAP:RNA (7.5 μM RNA) mixture produced PI positive droplets, however with no significant ThS signal (Figure S4), possibly as a result of a different type of LLPS permitting interactions. Whereas the binding of numerous NCAP monomers to a single RNA molecule at high protein to RNA ratio may promote amyloid fibril formation, a higher RNA to protein ratio may lead to a complex network of RNA-protein, RNA-RNA and protein-protein multivalent interactions(*8*), promoting phase separation, and suppressing amyloid formation.

**Fig. 4.**
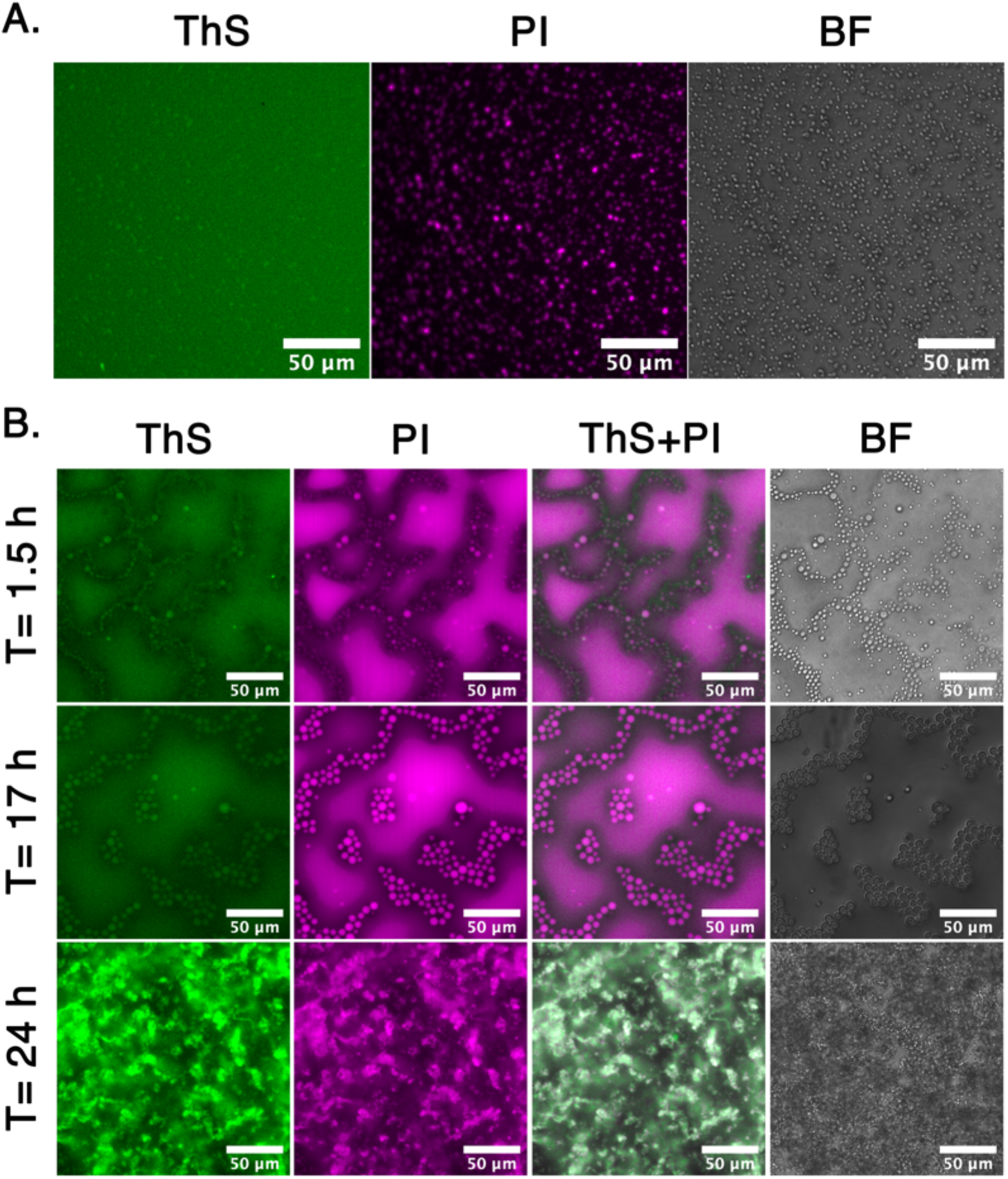
The NCAP and its LCD condense RNA into a separate liquid phase via a mechanism of amyloid-like aggregation as visualized by fluorescence microscopy. (A) LLPS and amyloid formed by 30 μM of NCAP incubated with 0.75 μM of Site2hp RNA (40:1 NCAP: Site2hp molar ratio). The NCAP droplets (~ 2-5 μm in diameter) exhibited RNA condensation activity (PI channel), and amyloid formation (ThS channel) during 17 h of incubation in 37 °C. Lack of RNA allowed no LLPS, while 7.5 μM of Site2hp (4:1 respective ratio) resulted in PI positive LLPS, but with no significant amyloid formation after 1.5 h and 17 h of incubation (Figure 4S). (B) LLPS and amyloid formed by 30 μM of the LCD when incubated with 7.5μM of Site2hp RNA. The LCD droplets exhibited RNA condensation activity (PI channel), and amyloid formation (ThS channel) that was enhanced in extended incubation periods. Upon 24 h of incubation, irregular shapes, seemingly of a solid-like hydrogel form, appeared.

The LCD segment, as opposed to the full length NCAP, form fibrils in lower protein:RNA ratios. Whereas we observed no LLPS or amyloid formation of the LCD in the 40:1 protein: RNA (0.75 μM) molar ratio, ThS and PI positive LLPS droplets of roughly 3-10 μm in diameter appeared in the 4:1 (7.5 μM RNA) molar ratio mixture during 17 h of incubation (Figure 4B & S4). 24 h of incubation led to the formation of gel-like particles of intense ThS signal (Figure 4B). This pronounced fibril formation of LCD at lower protein:RNA ratios may result when each LCD interacts and bridges between several RNA molecules, while the flexible LCD monomers allow amyloid-promoting LCD-LCD interactions to form a network of RNA and fibrils. We conclude that the protein to RNA ratio governs NCAP amyloid formation in the LLPS droplets.

To acquire the atomic structures necessary for the design of inhibitors of the amyloid form of NCAP and perhaps of its function, we identified and determined structures of its amyloid driving segments. Amyloid fibrils are stabilized by pairs of tightly mating ß-sheets, often with zipper-like interfaces. The sequence motifs that drive fibril-formation are termed steric zippers and can be identified by a computer algorithm(*24*). We identified three such steric zipper-forming 6-residue segments in the sequence of the LCD: _179_GSQASS_184, 217_AALALL_222_, and _243_GQTVTK_248_. X-ray structures confirmed that each segment forms amyloid-like fibrils within its crystal, stabilized by steric-zipper interfaces (Figure 5, S5 & Table 1). Both the segments GSQASS and GQTVTK form parallel, in-register β-sheets. The AALALL segment crystalized in two forms, both with antiparallel β-sheets (*25*). The weaker zipper interface of the second form incorporates polyethylene glycol (Figure S5), and we do not consider it further. Structural stability statistics of all four crystal structures are given in Table S1.

**Fig. 5.**
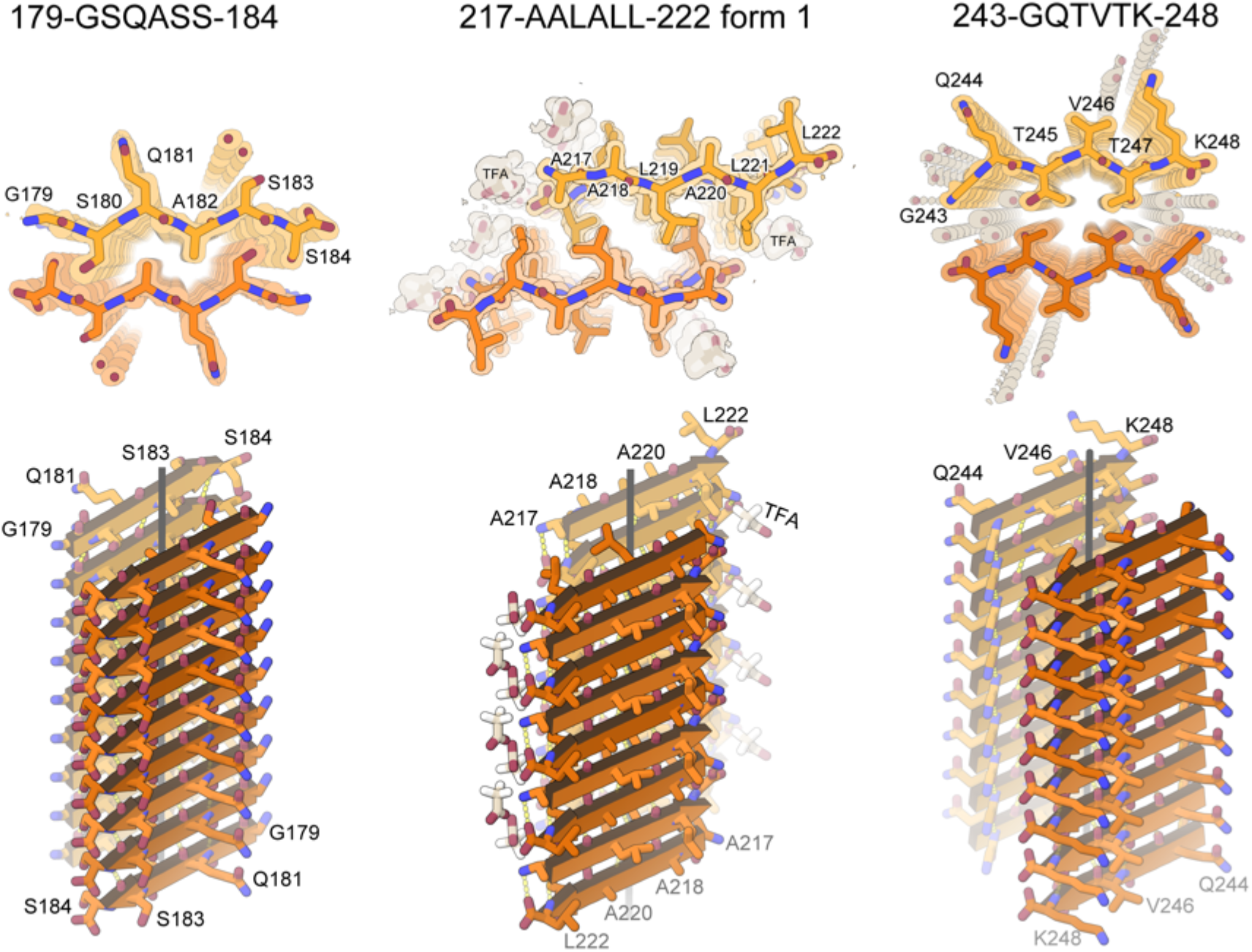
Amyloid-like association of NCAP segments revealed by crystallography. The upper row shows the quality of the fit of each model to its corresponding simulated annealing composite omit maps. The maps are contoured at the 1.0 sigma level. All structural features are well defined by the density. The view is directed down the fibril axis (crystal needle axis). Each chain shown here corresponds to one strand in a beta-sheet. Thousands of identical strands stack above and below the plane of the page making ~100 micron-long beta-sheets. The face of each beta-sheet of AALALL (PDB IDs: 7LTU, form 1) is symmetric with its back. However, GSQASS (PDB ID: 7LV2) and GQTVTK (PDB ID: 7LUZ) each reveal two distinct sheet-sheet interfaces: face-to-face and back-to-back. The tighter associated pair of sheets is shown here. The lower row shows 18 strands from each of the steric zippers at a view nearly perpendicular to the fibril axis. GSQASS and GQTVTK are parallel, in register sheets, mated with class 1 zipper symmetry. The AALALL zippers are antiparallel, in register sheets, mated with class 7 zipper symmetry. Trifluoroacetic acid (TFA) appear in the AALALL-form 1 steric zipper, and polyethylene glycol (PEG) binds form 2 (PDB ID: 7LUX; Figure S5).

**Table 1.** Crystallographic data collection and refinement statistics from SARS-CoV-2 NCAP segment

| Segment | <sup>179</sup>GSQASS<sup>184</sup> | <sup>217</sup>AALALL<sup>222</sup><br>Form 1 | <sup>217</sup>AALALL<sup>222</sup><br>Form 2 | <sup>243</sup>GQTVTK<sup>248</sup> |
| --- | --- | --- | --- | --- |
| **Data Collection** |  |  |  |  |
| Beamline | APS 24-ID-E | APS 24-ID-E | APS 24-ID-E | APS 24-ID-E |
| Space group | P2<sub>1</sub>2<sub>1</sub>2<sub>1</sub> | P1 | P2<sub>1</sub>2<sub>1</sub>2 | P2<sub>1</sub> |
| Resolution (Å) | 1.30 (1.39-1.30)* | 1.12 (1.18-1.12) | 1.30 (1.36-1.30) | 1.10 (1.17-1.10) |
| Unit cell dimensions: a,b,c (Å) | 4.77,13.60, 42.44 | 9.45, 11.34, 20.27 | 44.46, 9.54, 10.95 | 19.57, 4.78, 22.03 |
| Unit cell angles: α,β,γ (°) | 90.0,90.0,90.0 | 74.9, 79.1, 67.8 | 90.0, 90.0, 90.0 | 90.0, 94.0, 90.0 |
| Measured reflections | 1833 (338) | 5371 (323) | 4666 (550) | 4677 (344) |
| Unique reflections | 809 (139) | 2270 (136) | 1234 (139) | 1726 (170) |
| Overall completeness (%) | 93.2 (95.9) | 78.4 (31.1) | 93.0 (84.8) | 87.1 (50.9) |
| Overall redundancy | 2.3 (2.4) | 2.4 (2.4) | 3.8 (4.0) | 2.7 (2.0) |
| Overall R<sub>merge</sub> | 0.126 (1.04) | 0.084 (0.397) | 0.105 (0.808) | 0.085 (0.446) |
| CC<sub>1/2</sub> | 99.7 (56.7) | 98.5 (89.2) | 99.7 (54.4) | 99.5 (84.3) |
| Overall I/σ | 3.5 (0.7) | 5.9 (2.0) | 5.9 (1.8) | 6.0 (1.4) |
| **Refinement** |  |  |  |  |
| R<sub>work</sub> / R<sub>free</sub> | 0.259 / 0.253 | 0.158 / 0.197 | 0.217 / 0.248 | 0.133 / 0.177 |
| RMSD bond length (Å) | 0.015 | 0.009 | 0.010 | 0.009 |
| RMSD angle (°) | 1.4 | 1.3 | 1.6 | 1.5 |
| Number of peptide atoms | 40 | 180** | 40 | 93** |
| Number of water atoms | 2 | 1 | 1 | 12 |
| Number of other solvent atoms | 0 | 21 | 14 | 0 |
| Average B-factor of peptide (Å<sup>2</sup>) | 12.3 | 12.3 | 14.2 | 8.2 |
| Average B-factor of water (Å<sup>2</sup>) | 19.9 | 12.8 | 26.6 | 24.7 |
| Average B-factor other solvent (Å<sup>2</sup>) | N/A | 20.8 | 27.3 | N/A |
| PDB ID code | 7LV2 | 7LTU | 7LUX | 7LUZ |

\*Numbers in parentheses report statistics in the highest resolution shell.
\*\*Count includes hydrogen atoms.

Using our steric-zipper structures as templates, we designed 60 peptide-based candidate inhibitors aimed to cap the tips of our steric-zipper structures. We have found capping inhibitors to be effective in inhibiting aggregation and prion-like seeding of amyloid-forming proteins *(e.g.*(*26–32*)). To design NCAP fibrillation inhibitors, we implemented two approaches: Sequence/Structure-based design and Rosetta-based modeling(*33*). Both approaches identify bulky residues that block fibril extension by sterically interfering with the addition of another protein molecule to the fibril tip. Our Sequence/Structure-based approach maintains high sequence identity with the parent steric-zipper forming segment in order to exploit the homomeric tendency of amyloid aggregation, whereas the Rosetta-based approach explores sequence space more exhaustively (Figure S6). These approaches yielded 60 inhibitor candidates (Table S2), of which four (S17, S18, P22, and P29) showed the most pronounced inhibition of *in vitro* Site2hp-stimulated fibril formation of the LCD (Figure 6&S6). Candidates S18 and P22 exhibited dose dependent attenuation of LCD fibrillation, whereas P29 demonstrated significant, but equivalent, levels of inhibition at all tested concentrations. S17 showed increased attenuation of fibrillation in 1:2 and 1:1 S17:LCD molar ratios. Inhibition in the 2:1 S17:LCD molar ratio appears poorer, but this may be a false indication resulting from an underlying undetected aggregation of the inhibitor itself (Figure 6). When testing our lead compounds on the Site2hp-stimulated fibrillation of the NCAP, a different modulation pattern was revealed. The only compound showing some inhibition in 2:1 inhibitor:protein molar ratio was S18, whereas P22 showed no modulation of fibrillation and S17, and P29 actually enhanced NCAP fibrillation (Figure 6). Nevertheless, any change of NCAP fibril formation promoted by those compounds may conceivably be detrimental to the function of the virus.

**Fig. 6.**
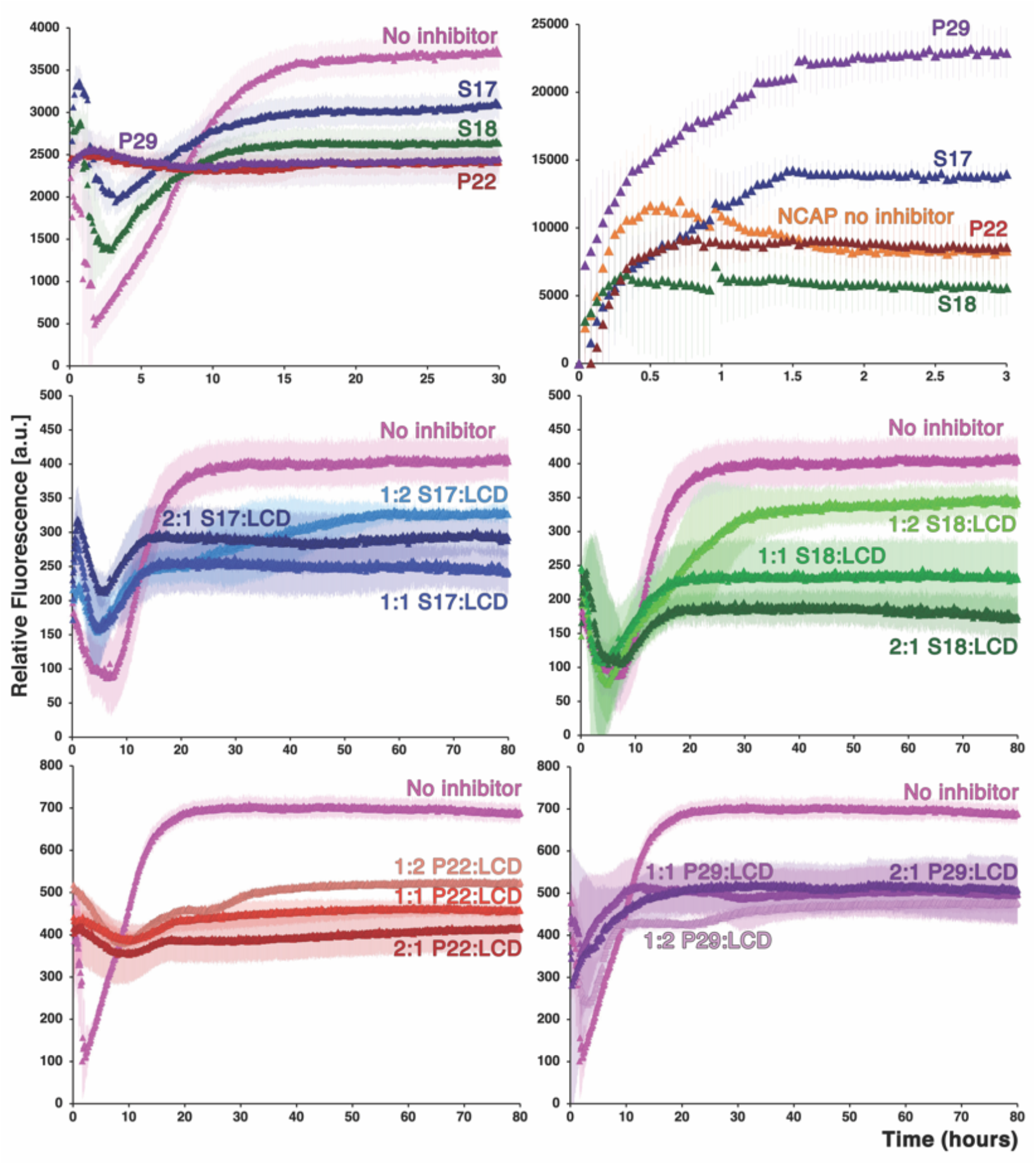
Inhibition of LCD and NCAP fibrillation observed *in vitro* using a ThT assay. Top: Best inhibitor candidates (S17, S18, P22, and P29; Figure S6, Table S2) reduce ThT fluorescence relative to a no-inhibitor control when individually mixed with LCD and site2hp RNA in a molar ratio of 8 inhibitor:4 LCD:1 RNA (left). The effect of those candidates on the fibrillation of the NCAP protein under the same conditions (right) is different, and suggestive of additional fibrillation prone regions in the protein. Bottom panels: Dose dependent inhibition evaluation of S17 (middle left), S18 (middle right), P22 (bottom left) and P29 (bottom right) on the LCD segment when mixed in 1:2, 1:1 or 2:1 inhibitor:protein and 4 protein: 1 site2hp RNA molar ratios. S17 shows signs of dose dependent inhibition, however, the 2:1 S17:LCD ratio mixture produces a higher fluorescence signal than 1:2 and 1:1. Aggregation of the inhibitor itself might account for this anomaly. P22 and S18, but not P29, are showing dose dependent inhibition. All experiments were repeated three times in triplicates, respective blanks (buffer+RNA+inhibitor) were subtracted, and representative curves are presented. Error bars represent SD.

## Discussion

Whereas amyloid fibrils were once considered irreversible aggregates, arising from the misfolding or denaturation of functional proteins, more recently scientists have discovered dozens of functional amyloid structures. These include, among others, components of bacterial biofilms and storage forms of human hormones(*34–39*). One subclass of these evolved amyloid fibrils are those formed by RNA binding proteins, such as FUS, TDP-43, and hnRNPA2, all of which function in RNA metabolism and are found in membraneless organelles such as stress granules (*10, 11, 20, 21, 12–19*). These proteins contain, in addition to RNA binding domains, low-complexity domains (LCDs) having sequences enriched in certain amino acid residues such as Glycine, Serine, Glutamine, and Tyrosine. Their LCDs, as well as the full proteins, have been found to undergo liquid–liquid phase separation, apparently as part of their cellular functions. Moreover, the LCDs have been found under conditions of high concentration to form amyloid-like fibrils and gels(*10, 14, 40*).

The NCAP protein of SARS-CoV-2 clearly belongs to this subclass of fibril-forming proteins. It contains both RNA-binding and low-complexity domains, it undergoes liquid–liquid phase separation(*1–8*), and as we show here, it forms amyloid-like fibrils. LLPS and fibril formation of NCAP, and its LCD, are stimulated by RNA, and the RNA-induced LLPS droplets contain amyloid fibrils, indicating that these processes are indeed correlated (Figures 3 & 4). These similarities suggest that NCAP fibrils function in its action of encapsulating RNA in viral replication. The hairpin-possessing viral RNA, Site2hp, exhibits strong binding to NCAP and some binding to its DD-C_term_ segment, but weak to no binding to N_term_-RBD and to the low-complexity domain itself (Figure 2 & S1). We interpret these results as indicating that the binding of RNA to NCAP involves several protein domains, as was found for the similar NCAP of SARS-CoV (Figure S7)(*23*). Together our observations suggest that NCAP is a fibril-forming protein and that its fibril formation is linked to its function of LLPS.

Evidence for the involvement of NCAP fibrils in virus function is strengthened by studies of other viruses. The NCAPs of Ebola and Measles, among others, also undergo LLPS as part of their viral life cycles. The LLPS of Measles’s NCAP was shown to mature from a liquid-like to a gel-like state, possibly as a mechanism to control phase separation(*41*). We similarly observed a transformation from a liquid-like to a gel-like state of the LCD segment upon extended incubation with Site2hp RNA (Figure 4). Also for human RNA-binding proteins, the formation and maturation of phase-separated droplets has been associated with increased fibril content(*12*).

If fibril formation is a feature of NCAP function, it follows that disruption of its fibril formation can disturb function. This possible disruption of SARS-CoV-2 function is the rationale of our design of inhibitors of its fibril formation. To design inhibitors, we needed atomic structures of the fibril-driving segments, and so we determined structures of the three predicted fibril-driving segments of the LCD (Figure 5). All three structures offer atomic-resolution information about these fibril-driving, “homo-steric-zipper” segments. Homo-steric zippers are formed from identical mating sheets. That is, the two sequence segments that interdigitate to form the steric zipper have the same sequence. In contrast, in hetero-steric zippers the mating sheets are different, and the two sequence segments that interdigitate have different sequences. In the near-atomic resolution cryoEM structures of amyloid fibrils (e.g. (*42, 43*)), both homo- and hetero-zippers are seen. Homo-zippers frequently stabilize the interaction of two identical protofilaments within a fibril. Hetero-zippers frequently stabilize the folds of individual protofilaments. In some cases, the stabilizing segment of a hetero-zipper of one of the sheets has also been found as a homozipper in a crystal structure. Of importance here, inhibitors designed based on template sequences of homo-zippers have been effective for inhibition of fibrils that contain that sequence in hetero-zippers(*26, 44*). In short, here we designed peptide inhibitors of the presently unknown structure of NCAP fibrils based on homo-zippers of its putative fibril-driving segments, with the reasonable expectation that these would inhibit fibril formation of LCD. Which they do.

To ensure success, armed with the structures of the three homo-zippers, we designed and tested 60 peptide-based candidates to interfere with fibril formation of NCAP (Figure S6 & Table S2). Inhibitors designed against the AALALL segment are the most efficient in attenuating in vitro LCD fibril formation, some in a dose dependent manner (Figure 6). The AALALL segment as the main driver of fibrils is consistent with our estimate that it forms the strongest stabilizing interactions (Table S1, ΔG° column). We note that our most effective candidates chosen according to their attenuation of the fibril formation of the LCD, do not exert the same inhibition on fibril formation of full NCAP. Some even enhanced fibril formation of the NCAP (Figure 6). This implies that additional segments of NCAP, such as the shorter low-complexity sequence near the C-terminus, are capable of driving NCAP fibrillation. It is possible, however, that candidates lacking in vitro inhibition of fibril formation of the full NCAP may nevertheless affect viral function. It will be necessary to evaluate all inhibitor candidates for their ability to reduce viral replication in SARS-CoV-2 infected cells.

Offering hope that designed inhibitors can disrupt function of various coronavirus strains, the sequences of our three steric-zipper forming segments are all moderately to highly conserved among homologous coronaviral NCAPs. The GQTVTK segment is the most conserved (Figure S7A). In fact, the entire LCD sequence of SARS-CoV-2 NCAP is very similar to that of the middle east respiratory syndrome (MERS) virus, and almost identical to that of SARS-CoV (Figure S7B). This high level of conservation points to evolutionary pressure to preserve the LCD sequence. The low genetic variation of viral NCAPs(*23*), may retard the development of viral resistance to our designed inhibitors. Furthermore, because our sequence/structure-based inhibitors were designed for sequence compatibility with the interacting interface in the amyloid spine, mutations arising as a drug resistance mechanism would be expected to impair protein function. That is, by aiming our inhibitors against conserved as well as non-conserved regions in the NCAP, it may be possible to provide both global and strain specific coronavirus therapeutics.

In summary, the SARS-CoV-2 NCAP protein undergoes liquid–liquid phase separation accompanied by formation of amyloid-like fibrils driven at least in part by the self-association of its low-complexity domain and interaction with RNA. Based on three atomic structures of the fibril-driving segments of NCAP, we designed peptides that affect NCAP fibril formation. These peptides may offer a new approach to anti COVID therapies. Experience shows that even efficient vaccines rarely eradicate viral diseases and their legacies of morbidity and mortality(*45*), so therapies are needed. In the case of COVID-19, lingering symptoms(*46*) are not uncommon, including neurological complications(*47*). Because amyloid fibrils are associated with numerous dementias and movement disorders(*22, 48*) management of fibril formation of NCAP may have particular importance.

## Materials and Methods

### Molecular Biology Reagents

Phusion HF DNA polymerase, Quick Ligase and restriction enzymes were obtained from New England BioLabs. Custom oligonucleotides were synthesized by IDT (Coralville, IA).

### Low Complexity Domain Prediction

Amino acid sequences in the proteome of SARS-CoV-2 were evaluated for low complexity using SEG(*49*) with default settings: window length = 12, trigger complexity 2.2, extension complexity 2.5. A sequence was determined to be a low complexity domain if it contained at least 25 residues scored as low complexity with at most 5 interrupting non-low complexity residues(*49*).

#### Construct design

Full length SARS-CoV-2 nucleoprotein gene and its fragments were PCR amplified from 2019-nCoV Control Plasmid (IDT Inc, Cat No 10006625) and spliced with N-terminal 6xHis-SUMO tag(*50*) using splicing by overlap extension (SOE) technique(*51*). 5’ KpnI and 3’ SacI restriction sites introduced with the flanking primers were used to ligate resulting constructs into pET28a vector. When needed, additional round of SOE was performed to generate internal nucleoprotein deletion mutants. Construct sequences were confirmed by Sanger sequencing (Laragen, Culver City, CA).

#### Protein Expression & Purification

NCAP constructs were expressed as fusions to 6xHis-SUMO, linked via an N-terminal Ulp1 cleavage site (6xHis-SUMO-Ulp1site-NCAP). Plasmids were transformed into *Escherichia coli* Rosetta2 (DE3) strain (MilliporeSigma cat. no 71-397-4) and small-scale cultures were grown at 37 °C overnight in LB with 35 μg/mL kanamycin and 25 μg/mL chloramphenicol. TB with 35 μg/μL kanamycin was inoculated with overnight starter culture at a 1:100 ratio and large-scale cultures were grown at 37 °C, 225 rpm until the OD600 reached ~0.6. Protein expression was induced with 1 mM IPTG and cultures were further incubated with shaking at 28 °C overnight, then harvested at 5000 *g* at 4 °C for 15 minutes. Bacterial pellets were either used right away or stored at −20 °C. Pellets were re-suspended in ~200 mL chilled Buffer A (20 mM Tris pH 8.0, 1 M NaCl), supplemented with Halt Protease Inhibitor Cocktail (ThermoScientific cat. no 87785), per 2-4 L culture, and sonicated on ice at 80 % amplitude for a total sonication time of 15 minutes, with regular pauses to not exceed 15 °C. Cell debris was removed via centrifugation at 24,000 *g* at 4 °C for 30-60 minutes, then filtered twice through 0.45-*μm* high particulate syringe filters (MilliporeSigma cat. no SLCRM25NS), and imidazole added to 5mM. Filtered clarified lysate was loaded onto HisTrap HP columns (GE Healthcare) and proteins eluted over a step gradient with Buffer B (20 mM Tris pH 8.0, 1 M NaCl, 500 mM imidazole), with extensive low imidazole (<20 %) washes to improve purity. NCAP proteins generally eluted in 20-50 % Buffer B, and fractions were analyzed by SDS-PAGE, pooled and dialyzed against 20 mM Tris pH 8.0, 250 mM NaCl at 4 °C overnight. Following dialysis, the sample was concentrated using Amicon Ultra-Centrifugal filters (MilliporeSigma) and urea added up to 1 M final concentration if precipitation was observed. Ulp1 protease (homemade) was added at a 1:100-1:200 w/w ratio to protein, along with 1 mM DTT, and the sample was incubated at 30 °C, 195 rpm for 1-2 hours. Following cleavage, NaCl was added to the sample to 1M final concentration to reduce aggregation, and the sample was incubated with HisPur Ni-NTA resin (ThermoScientific cat. no PI88222) equilibrated in Buffer A at 25 °C, 140 rpm for 30 minutes. Cleaved NCAP protein was separated from resin via gravity flow column, and resin was washed twice with Buffer A then twice with Buffer A + 5 mM imidazole and finally with Buffer B. Flow-thru and appropriate washes were concentrated and flash frozen for storage or further purified by gel filtration. Directly prior to gel filtration, the sample was centrifuged at 21,000 *g* for 30 minutes at 4 °C to remove large aggregates. Soluble protein was injected on a HiLoad Sephadex 16/600 S200 (for constructs larger than ~25 kDa) or S75 (for constructs smaller than ~25 kDa) (GE Healthcare) equilibrated in SEC buffer (20 mM Tris pH 8.0, 300 mM NaCl) and run at a flow rate of 1 mL/min. Elution fractions corresponding to monomeric protein were assessed by SDS-PAGE for purity, and confirmed to have low RNA contamination as assessed by 260/280 nm absorbance ratio. Pooled fractions were concentrated and 0.2-*μm* filtered. Protein concentration was measured by A280 absorbance using a NanoDrop One (ThermoScientific) and the calculated extinction coefficient, and aliquots were flash frozen and stored at −80°C.

#### RNA Synthesis and Purification

The nucleic acid sequence corresponding to Site2hp was cloned from a gBlock (IDT) of the first 1000 nucleotides of the 5’-end of the SARS-CoV-2 genome into pUC19 vectors using the restriction sites EcoRI and KpnI. The clone was sequence-confirmed and the miniprep was used as a template for PCR. The forward primer for PCR containing the T7 promoter sequence was biotinylated on the 5’ end for removal of PCR template after transcription. The PCR product was purified by HiTrap Q HP column. The running buffer solutions (0.2-*μm* filtered) contained 2 M NaCl, 10 mM HEPES pH 7.0, and 10 mM NaCl, 10 mM HEPES pH 7.0. The purified PCR products were concentrated using Amicon Ultra centrifugal filter units (Millipore) and buffer exchanged against 10 mM HEPES pH 7.0. Transcription reactions ranging from 5 to 30 mL were set up. Each transcription reaction was incubated at 37°C with gentle shaking for one hour. After transcription, streptavidin beads (ThermoFisher) were added to the transcription and set on a rotator at room temperatures for an additional 15 minutes. The transcription was then centrifuged at 500xg for 10 minutes at 4°C. The supernatant was decanted and the remaining pellet containing any PCR template remaining was discarded. The transcription was then purified by 5-mL HiTrap Q HP column in several rounds, each loading around 5 mL of the transcription into the column. The purified RNA was concentrated using Amicon Ultra-15 (Millipore) and buffer exchanged with 10 mM HEPES pH 7.0.

### RNA binding assays

#### Size Exclusion Chromatography (SEC)

SEC experiments were conducted at room temperature. All samples were kept on ice and briefly centrifuged before being run through the Superdex 200 Increase 5/150 GL column. The running buffer for all the constructs was 1xPBS. All protein-only runs contained 24 μM protein in 1xPBS, and RNA-only runs contained 12 μM RNA in 1xPBS. 24 μM of protein and 12 μM of Site2hp RNA were used for protein-RNA mixtures, free protein and free RNA mixtures included only the appropriate component in the same concentration mentioned. Binding was determined based on the shift from the RNA only elusion peak to the protein-RNA mixture elusion peak. The molar ratio of protein:RNA in the mixtures was calculated using the following equation (*52*):

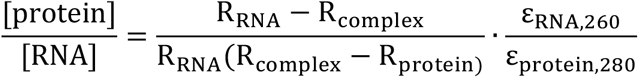

#### Native gel analysis

The interactions between NCAP proteins and Site2hp were analyzed using 8% native polyacrylamide gels, which were run in 1xTBE buffer at room temperature. The protein-only mixtures were 48 μM and the Site2hp-only mixtures were 12 μM. The protein and RNA were mixed in a 2:1 ratio (24 μM protein, 12 μM Site2hp) and in a 4:1 ratio (48 μM protein, 12 μM Site 2hp). The protein-RNA mixtures were mixed well and incubated at room temperature for ~40 min. After the incubation, the samples were diluted twelve-fold in 1xPBS to a final concentration of 1 μM Site2hp for better visualization on the gel. The gels were stained using Sybr Green II dye and imaged using the Fotodyne, 2MPix monochrome Analyst^®^ Express system. The images were analyzed using Quantity One (BIO-RAD).

#### Peptide Synthesis and Purification

The NCAP segments 179-GSQASS-184 and 243-GQTVTK-248 were synthesized and purified by LifeTein company.

The NCAP segment 217-AALALL-222 was synthesized and purified in-house as H-AALALL-OH. Peptide synthesis was carried out at 0.1 mmol scale. A 2-chlorotrityl chloride resin (Advanced Chemtech) was selected as the solid support with a nominal loading of 1.0 mmol/g. Each loading of the first amino acid was executed by adding 0.1 mmol of Fmoc-Leu-OH (Advanced Chemtech FL2350/ 32771) and 0.4 mmol of diisopropylethylamine (DIPEA), dissolved in 10 mL of dichloromethane (DCM), to 0.5 grams of resin. This mixture was gently agitated by bubbling with air. After 30 minutes, the supernatant was drained, and the resin was rinsed twice with 15 mL aliquots of capping solution, consisting of 17:2:1 DCM/MeOH/DIPEA. With the first amino acid loaded, the elongation of each polypeptide was completed in a CEM Liberty Blue™ Microwave Peptide Synthesizer. A 1.0 M solution of N,N’-diisopropylcarbodiimide (DIC) in DMF was used as the primary activator, and a 1.0 M solution of ethyl cyanohydroxyiminoacetate (oxyma) in DMF, buffered by 0.1 M of DIPEA was used as a coupling additive. The Fmoc-L-Ala-OH used was also purchased from Advanced Chemtech (FA2100/ 32786). The microwave synthesizer utilizes 0.2 M solutions of each amino acid. For the deprotection of N-termini, Fmoc protecting groups, a 9% w/v solution of piperazine in 9:1 N-Methyl-2-Pyrrolidone to EtOH, buffered with 0.1 M of oxyma was used. For 0.1 mmol deprotection reactions, 4 mL of the above deprotection solution was added to the resin. The mixture was then heated to 90°C for 2 minutes while bubbled with nitrogen gas. The solution was drained, and the resin washed with 4 times with 4 mL aliquots of DMF. For 0.1 mmol couplings, 2.5 mL of 0.2 M amino acid solution (0.5 mmol) was added to the resin along with 1 mL of the DIC solution (1.0 mmol) and 0.5 mL of oxyma solution (0.5 mmol). This mixture was agitated by bubbling for 2 minutes at 25 °C and then heated to 50 °C followed by 8 minutes of bubbling. After the last deprotection, the resin was washed with methanol, diethyl ether, dried over vacuum, and introduced to a cleavage cocktail consisting of: 20 mL of trifluoroacetic acid (TFA); 0.50 mL of water; 0.50 mL of triisopropylsilane (TIS). After 2 hours of vigorous stirring, the mixture was filtered, and the filtrate concentrated in vacuo. The residue was triturated with cold diethyl ether, and precipitated, crude peptide was collected by filtration. The crude peptide was then purified by RP-HPLC, using an Interchim puriFlashR 4125 Preparative Liquid Chromatography System equipped with a Luna (Phenomenex, C18(2), 5 μm, 100 Å, 30 x 100 mm) column. For purification, two buffer systems were utilized. Initial purifications and salt exchanges were executed with a 13 mM aqueous solution of trifluoroacetic acid (TFA; [A]) and a 2:3 water to acetonitrile solution, buffered by 13 mM of TFA ([B]). For better resolution of diastereomers and other impurities, ultrapure water, buffered by 14 mM of HClO4, and a 2:3 water to acetonitrile solution, buffered by 5.6 mM of HClO4, were selected as mobile phases A and B, respectively. The purity of the purified fractions was analyzed by RP-HPLC, using an Agilent 1100 Liquid Chromatography System equipped with a Kinetex (Phenomenex, C18, 5 μm, 100 Å, 4.6 x 250 mm) column. Ultrapure water with 0.1% TFA, and a 1:9 water to acetonitrile solution with 0.095% TFA were selected as mobile phases [A] and [B], respectively. The flow rate was set at 1.0 mL/min and the gradient used is detailed in Table S4. The UV absorption at 214 nm was monitored. The resulting chromatogram is shown in Figure S8.

#### Crystallization of NCAP segments

The N-protein segment 217-AALALL-222 crystallized in batch just before the purification by RP-HPLC. The peptide had been deprotected and cleaved from the resin, triturated with cold diethyl ether, and precipitated. Most of the product had been collected via filtration, but some residual peptide remained in the round bottom flask and we intended to use this residual to check the peptide purity by analytical HPLC. We dissolved this residual with water, acetonitrile, and TFA in a ratio of approximately 45:45:10 and transferred to a 1 mL glass vial for HPLC injection. The solution was left in the sample holder and crystals were observed to form after a week. These crystals were needle-shaped. Some of these crystals were retained for crystal structure determination. The bulk of the peptide was further purified, as described above. The purified peptide was dissolved at 10 mg/mL concentration in 19.6 mM LiOH. Crystals grew by the hanging drop vapor diffusion method. The UCLA Crystallization Facility set up crystallization trays with a Mosquito robot dispensing 200 nL drops. Needle shaped crystals grew at 20 °C using a reservoir solution composed of 30 % (w/v) polyethylene glycol (PEG) 3000 and 0.1 M n-Cyclohexyl-2-aminoethanesulfonic acid (CHES), pH 9.5. The purified NCAP segment 179-GSQASS-184 was dissolved in water at 100 mg/mL concentration. Hanging drop crystallization trays were set using 200 nL drops. Needle shaped crystals grew at 20 °C using a reservoir solution composed of 1.0 M Na,K tartrate, 0.2 M Li_2_SO_4_, and Tris pH 7.0. Needle-shaped crystals appeared immediately after setting up the tray. The purified NCAP segment 243-GQTVTK-248 was dissolved in water at 68 mg/mL concentration. Hanging drop crystallization trays were set using 200 nL drops. Needle-shaped crystals appeared within 1 day at 20 °C using a reservoir solution composed of 2.0 M (NH4)2SO4, 0.1 M sodium HEPES, pH 7.5, and 2 % (v/v) PEG 400.

#### Crystallographic Structure Determination of NCAP segments

Microfocus X-ray beam optics were required to measure crystal diffraction intensities from our crystals since they were needle-shaped, and less than 5 microns thick. We used microfocus beamline 24-ID-E of the Advanced Photon Source located at Argonne National Laboratory. Crystals were cooled to a temperature of 100 K. Diffraction data were indexed, integrated, scaled, and merged using the programs XDS and XSCALE(*53*). Data collection statistics are reported in Table 1. Initial phases for AALALL and GSQASS were obtained by molecular replacement with the program Phaser(*54*) using a search model consisting of an ideal beta-strand with sequence AAAAAA. Phases for GQTVTK were obtained by direct methods using the program ShelxD(*55*). Refinement was performed using the program Refmac(*56*). Model building was performed using the graphics program Coot(*57*). Structure illustrations were created using PyMOL (*58*).

#### Rosetta-based peptide inhibitor design

Crystal structures of LCD segments GSQASS, GQTVTK and AALALL with TFA were used as templates for the design of peptide inhibitors in Rosetta3 software(*33*). 5 layers of the steric zipper structure was generated. A 6-residue L-peptide chain was placed at the top or bottom of the fibril-like structure. Rosetta Design was used to sample all amino acids and their rotamers on the sidechains of the fixed peptide backbone. The lowest energy conformations of the sidechains were determined by minimizing an energy function containing terms for Lennard-Jones potential, orientation-dependent hydrogen bond potential, solvation energy, amino acid-dependent reference energies, and statistical torsional potential dependent on backbone and sidechain dihedral angles. Buried surface area and shape complementarity were scored by AREAIMOL(*59*) and Sc(*60*), respectively, from the CCP4 suite of crystallographic programs(*61*). Design candidates were selected based on their calculated binding energy to the top or bottom of the fibril-like structure, shape complementarity, and propensity for self-aggregation. The structural model of each candidate peptide was manually inspected in PyMOL(*58*). Many computational designs produced sequences with high hydrophobic content, thus two arginine residues were added onto the C-terminal end to increase peptide solubility.

### Thioflavin-T fibrillation kinetic assay

All solutions in this experiment were prepared using DNase/RNase-free water (ultrapure water), and filtered twice using a 0.22-*μm* syringe filter. Preparations were done under sterile conditions and using filter pipette tips to ensure RNA preservation. Thioflavin T (ThT) stock solution was freshly prepared from powder (Sigma, CAS ID: 2390-54-7) at a concentration of 20mM in DNase/RNase ultrapure water, followed by 0.22-*μm* filtration. The NCAP protein and its segments were diluted into 235 μM working stocks using 20 mM Tris, 300 mM NaCl, pH 8.0 buffer. The Site2hp RNA was diluted into 75 μM working stock using 10 mM HEPES, pH 7.0 buffer. Each reaction sample contained 300 μM ThT, 30 μM protein and 7.5 μM RNA (for samples containing RNA), in 1xPBS pH 7.4. Blank samples contained everything but the protein. The reaction was carried out in a black 384-well clear-bottom plate (NUNC 384), covered with an optical film (Corning Sealing Tape Universal Optical) and incubated in a plate reader (BMG LABTECH FLUOstar Omega), at 37 °C, with 700 rpm double orbital shaking for 30 seconds before each measurement. ThT fluorescence was measured with excitation and emission wavelengths of 430 and 485 nm, respectively, and measurements were taken every 5 minutes for up to 1000 cycles. Measurements were made in triplicates for each sample. All triplicate values were averages, and blank readings from samples without protein are averaged and subtracted from the appropriate curves. The results were plotted against time with calculated standard deviation (SD) as error bars. The entire experiment was repeated at least three times on different days.

#### Evaluation of peptide-based inhibitor candidates

All solutions in this experiment were prepared using DNase/RNase-free water (ultrapure water), and filtered twice using a 0.22-*μm* syringe filter. Preparations were done under sterile conditions and using filter pipette tips to ensure RNA preservation. peptide-based Inhibitor candidates were synthesized by GenScript and LifeTein. Lyophilized peptide powders were separately dissolved to 10 mM concentration in DMSO or water, and stored at −20 °C. Peptides were added in 2:1, 1:1, or 1:2 protein:inhibitor molar ratio (as depicted in Figures 5 & S6) to 30 μM of either the NCAP protein or its LCD segment, and complemented with 7.5 μM of Site2hp RNA (4:1 protein:RNA molar ratio) in PBS buffer in a 384 black-well clear-bottom plate (Nunc). A ThT stock was prepared as described above and added to the well to 300 μM final concentration. The plates were immediately covered with an optical film (Corning Sealing Tape Universal Optical) and a ThT fibrillation kinetic assay was performed as described above to evaluate the fibrillation of the NCAP and LCD proteins in the absence and presence of different concentrations of peptide-inhibitor candidates.

#### Turbidity kinetic absorbance measurement assay

All solutions in this experiment were prepared using DNase/RNase-free water (ultrapure water), and filtered twice using a *0.22-μm* syringe filter. Preparations were done under sterile conditions and using filter pipette tips to ensure RNA preservation. Protein and RNA working solutions were prepared as described above for the ThT (no inhibitor) experiments. Each reaction sample contained 30 μM protein and 7.5 μM RNA (for samples containing RNA), in 1xPBS pH 7.4. Blank samples contained everything but the protein. The reaction was carried out in a black 384-well clear-bottom plate (NUNC 384), covered with an optical film (Corning Sealing Tape Universal Optical) and incubated in a plate reader (BMG LABTECH FLUOstar Omega), at 37 °C, with mixing for 5 seconds before measurements and 200 seconds between measurements. Turbidity was measured with absorbance (OD) at 600nm, and measurements were taken every 5 minutes for up to 14 hours. Measurements were made in triplicates for each sample. All triplicate values are averages, and appropriate blank readings (from samples without the protein) are averaged and subtracted from all other readings. The results were plotted against time with calculated SD as error bars. The entire experiment was repeated at least three times on different days.

#### Transmission electron microscopy (TEM)

NCAP and its segments were diluted into 235 μM working stocks using 20 mM Tris, 300mM NaCl, pH 8.0 buffer. The Site2hp RNA was diluted into 250 μM working stock using in 10 mM HEPES, pH 7.0 buffer. Each reaction sample contained 100 μM protein and 25 μM RNA (for samples containing RNA), in 1xPBS pH 7.4. A sample of only 25 μM Site2hp RNA was made as a control. The fibrillation reaction was carried out in parafilmed PCR tubes and incubated in a floor shaker (Torrey Pines Scientific Inc, Orbital mixing chilling/heating plate) at 37 °C, with highest mixing speed for 14 days. Samples (4 μL) were applied directly onto 400-mesh copper TEM grids with Formvar/Carbon support films (Ted Pella), glow discharged (PELCO easiGlowxs) for 45 seconds at 15 mA immediately before use. Grids were incubated with the samples for 2 minutes. Then, the samples were blotted off and the grids were washed three times with water and once with 2 % uranyl acetate solution with blotting in between. The grids were then incubated with 6 μL of uranyl acetate solution for 30 seconds before blotting. Micrographs were recorded using a FEI Tecnai T12 microscope at room temperature with an accelerating voltage of 120kV. Images were recorded digitally with a Gatan US 1000 CCD camera, using the Digital Micrograph^®^ software, and processed in the ImageJ^®^ software.

#### LLPS assay

All solutions in this experiment were prepared using DNase/RNase-free water (ultrapure water), and filtered twice using a 0.22-*μm* syringe filter. Preparations were done under sterile conditions and using filter pipette tips to ensure RNA preservation. The NCAP, LCD, N_term_-RBD, and DD-C_term_ protein stock solutions were centrifuged for 15 min, at 15,000 x g and 4 °C to remove large aggregates. The supernatant of each protein was separately diluted to 300 μM stock concentration using the same buffer of the original protein stock, containing 20 mM Tris, 300 mM NaCl, pH 8.0, followed by further dilution in 1xPBS to 50 μM working solutions. Site2hp RNA, stored in −20 °C, was thawed, then boiled at 95 °C for 3 min and placed immediately back on ice. The RNA was separately diluted into its stock buffer of 10 mM HEPES pH 7.0 to 187.5 μM and 18.75 μM stocks individually, then further diluted with 1xPBS to 25 μM and 2.5 μM stocks. Respective protein and RNA free solutions containing only the stock buffers were prepared as controls. Fresh Thioflavin S (ThS) solution was prepared from powder (originally from MP Biomedicals, CAS ID: 1326-12-1) in ultrapure water to 0.01 % (w/v) and filtered twice. A 1 mg/mL propidium iodide (PI) solution (Invitrogen, Cat# P3566) was diluted 10 folds in ultrapure water and filtered as well. The dyes were further mixed in 1xPBS to 0.002 % (w/v) ThS and 0.08 mg/mL PI. A single dye, and no dye controls were performed to ensure that under those experimental conditions the dye exert no effect on LLPS formation or fibrillation of the protein mixtures. Each protein, Site2hp RNA, and the dyes were mixed in wells of 384 black-well clear-bottom plate (Nunc) to final concentrations of 0.0002 % (w/v) ThS, 8 μg/mL PI, 30 μM of each individual protein or blank, and 0, 0.75, and 7.5 μM of Site2hp RNA to reach 1:0, 40:1 and 4:1 protein:RNA molar ratios respectively (as well as the appropriate protein free blanks). The plate was immediately covered with an optical film (Corning Sealing Tape Universal Optical). A control experiment in which the dyes were added to the samples 24 h post incubation in 37 °C was performed to ensure no dye effect on LLPS and fibrillation kinetics. Images were obtained using an ImageXpress XL (Molecular Devices, Sunnyvale) equipped with an 20x S Plan Fluor objective (NA 0.45) with Phase contrast and extra-long working distance (ELWD), environmental control module and transmitted light option with phase contrast light source. The imaging system was equilibrated to 37 °C for 2 hours before use and the plate was inserted as the measurement started. At least four sites were imaged for each well and each individual site was automatically focused using the IR laser autofocus of the ImageXpress. The plate was imaged in phase contrast channel with 2000ms exposure time, in the DAPI channel with 69 ms exposure time and in the Cy3 channel with 10ms exposure time. Measurements were taken at different time points as depicted in Figures 4 & S4. All images were taken at full camera resolution without any binning. The camera used was a Zyla scientific CMOS camera with 5.5 megapixel and a dual camera link interface. All images were recorded live into the MetaXpress Database and exported from there as need for downstream applications. Representative sites and time points images are presented in Figures 4 & S4. Note that although ThS has a high background signal, brighter foci are indicative of the dye increased local concentration resulting from its binding to amyloid fibrils.

## Acknowledgments

We thank Megan Bentzel, Jose Rodriguez, and Mark Arbing for discussions. We thank the staff at the Northeastern Collaborative Access Team, which is funded by the National Institute of General Medical Sciences from the National Institutes of Health (P30 GM124165). The Eiger 16M detector on the 24-ID-E beam line is funded by a NIH-ORIP HEI grant (S10OD021527). The Advanced Photon Source, a U.S. Department of Energy (DOE) Office of Science User Facility operated for the DOE Office of Science by Argonne National Laboratory under Contract No. DE-AC02-06CH11357.

## Author contribution

Author contributions: All authors contributed to conceptualization and to various aspects of the manuscript. Protein constructs and cloning: P.M.S, Luk.S. Protein preparation and experimentation: X.C., J.B., S.G, E.T.F, R.A, J.L, Y.J. RNA preparation and experimentation: C.T., Y.L. Peptide preparation: C.T.Z. X-ray crystallography: C.T.Z, M.R.S., M.B, J.L, K.H, G.F, D.C. Fluorescence and electron Microscopy: E.T.F., X.C., D.B. Computational analysis and inhibitor design: G.M.R., P.M.S., Y.J., K.A.M., E.T.F., Lor.S. Writing and figure preparation: E.T.F., M.R.S., F.G., D.S.E. Technical support: D.H.A, Project management: E.T.F., M.R.S., J.A.R., F.G., D.S.E.

## Data Availability

The atomic coordinates were deposited in the RCSB Protein Data Bank (PDB) under accession numbers 7LV2, 7LTU, 7LUX, and 7LUZ.All other data is available in the main and supplementary text and upon request.

## Supplementary Materials

Figures S1-S9; Tables S1-S4; Movies S1

## References

1. A. Savastano, A. Ibáñez de Opakua, M. Rankovic, M. Zweckstetter, Nucleocapsid protein of SARS-CoV-2 phase separates into RNA-rich polymerase-containing condensates. Nat. Commun. 11, 1–10 (2020).

2. H. Chen, Y. Cui, X. Han, W. Hu, M. Sun, Y. Zhang, P. H. Wang, G. Song, W. Chen, J. Lou, liquid–liquid phase separation by SARS-CoV-2 nucleocapsid protein and RNA. Cell Res. 30, 1143–1145 (2020).

3. J. Cubuk, J. Alston, J. J. Incicco, S. Singh, M. Stuchell-Brereton, M. Ward, M. Zimmerman, N. Vithani, D. Griffith, J. Wagoner, G. Bowman, K. Hall, A. Soranno, A. Holehouse, The SARS-CoV-2 nucleocapsid protein is dynamic, disordered, and phase separates with RNA. bioRxiv Prepr. Serv. Biol. (2020), doi:10.1101/2020.06.17.158121.

4. S. Lu, Q. Ye, D. Singh, E. Villa, D. W. Cleveland, K. D. Corbett, The SARS-CoV-2 Nucleocapsid phosphoprotein forms mutually exclusive condensates with RNA and the membrane-associated M protein. bioRxiv Prepr. Serv. Biol. (2020), doi:10.1101/2020.07.30.228023.

5. T. M. Perdikari, A. C. Murthy, V. H. Ryan, S. Watters, M. T. Naik, N. L. Fawzi, SARS-CoV-2 nucleocapsid protein phase-separates with RNA and with human hnRNPs. EMBO J. 39 (2020), doi:10.15252/embj.2020106478.

6. S. M. Cascarina, E. D. Ross, A proposed role for the SARS-CoV-2 nucleocapsid protein in the formation and regulation of biomolecular condensates. FASEB J. 34, 9832–9842 (2020).

7. C. Iserman, C. A. Roden, M. A. Boerneke, R. S. G. Sealfon, G. A. McLaughlin, I. Jungreis, E. J. Fritch, Y. J. Hou, J. Ekena, C. A. Weidmann, C. L. Theesfeld, M. Kellis, O. G. Troyanskaya, R. S. Baric, T. P. Sheahan, K. M. Weeks, A. S. Gladfelter, Genomic RNA Elements Drive Phase Separation of the SARS-CoV-2 Nucleocapsid. Mol. Cell. 80, 1078–1091.e6 (2020).

8. C. R. Carlson, J. B. Asfaha, C. M. Ghent, C. J. Howard, N. Hartooni, M. Safari, A. D. Frankel, D. O. Morgan, Phosphoregulation of Phase Separation by the SARS-CoV-2 N Protein Suggests a Biophysical Basis for its Dual Functions.Mol. Cell. 80, 1092–1103.e4 (2020).

9. Q. Ye, A. M. V. West, S. Silletti, K. D. Corbett, Architecture and self-assembly of the SARS-CoV-2 nucleocapsid protein. Protein Sci. 29, 1890–1901 (2020).

10. M. Kato, T. W. Han, S. Xie, K. Shi, X. Du, L. C. Wu, H. Mirzaei, E. J. Goldsmith, J. Longgood, J. Pei, N. V. Grishin, D. E. Frantz, J. W. Schneider, S. Chen, L. Li, M. R. Sawaya, D. Eisenberg, R. Tycko, S. L. McKnight, Cell-free Formation of RNA Granules: Low Complexity Sequence Domains Form Dynamic Fibers within Hydrogels. Cell. 149, 753–767 (2012).

11. S. Xiang, M. Kato, L. C. Wu, Y. Lin, M. Ding, Y. Zhang, Y. Yu, S. L. McKnight, The LC Domain of hnRNPA2 Adopts Similar Conformations in Hydrogel Polymers, Liquid-like Droplets, and Nuclei. Cell. 163, 829–839 (2015).

12. Y. Lin, D. S. W. Protter, M. K. Rosen, R. Parker, Formation and Maturation of Phase-Separated Liquid Droplets by RNA-Binding Proteins. Mol. Cell. 60, 208–219 (2015).

13. A. Molliex, J. Temirov, J. Lee, M. Coughlin, A. P. Kanagaraj, H. J. Kim, T. Mittag, J. P. Taylor, Phase Separation by Low Complexity Domains Promotes Stress Granule Assembly and Drives Pathological Fibrillization. Cell. 163, 123–133 (2015).

14. M. P. Hughes, M. R. Sawaya, D. R. Boyer, L. Goldschmidt, J. A. Rodriguez, D. Cascio, L. Chong, T. Gonen, D. S. Eisenberg, Atomic structures of low-complexity protein segments reveal kinked β sheets that assemble networks. Science. 359, 698–701 (2018).

15. E. W. Martin, T. Mittag, Relationship of Sequence and Phase Separation in Protein Low-Complexity Regions. Biochemistry. 57, 2478–2487 (2018).

16. B. A. Berning, A. K. Walker, The Pathobiology of TDP-43 C-Terminal Fragments in ALS and FTLD. Front. Neurosci. 13, 335 (2019).

17. W. M. Babinchak, R. Haider, B. K. Dumm, P. Sarkar, K. Surewicz, J.-K. Choi, W. K. Surewicz, The role of liquid–liquid phase separation in aggregation of the TDP-43 low-complexity domain. J. Biol. Chem. 294, 6306–6317 (2019).

18. S. Elbaum-Garfinkle, Matter over mind: Liquid phase separation and neurodegeneration. J. Biol. Chem. 294, 7160–7168 (2019).

19. T. M. Franzmann, S. Alberti, Prion-like low-complexity sequences: Key regulators of protein solubility and phase behavior. J. Biol. Chem. 294, 7128–7136 (2019).

20. J. Shorter, Phase separation of RNA-binding proteins in physiology and disease: An introduction to the JBC Reviews thematic series. J. Biol. Chem. 294, 7113–7114 (2019).

21. Y. Xing, A. Nandakumar, A. Kakinen, Y. Sun, T. P. Davis, P. C. Ke, F. Ding, Amyloid Aggregation under the Lens of Liquid–Liquid Phase Separation. J. Phys. Chem. Lett. 12, 368–378 (2020).

22. D. Eisenberg, M. Jucker, The amyloid state of proteins in human diseases. Cell. 148, 1188–1203 (2012).

23. C. K. Chang, M. H. Hou, C. F. Chang, C. D. Hsiao, T. H. Huang, The SARS coronavirus nucleocapsid protein – Forms and functions. Antiviral Res. 103, 39–50 (2014).

24. L. Goldschmidt, P. K. Teng, R. Riek, D. Eisenberg, Identifying the amylome, proteins capable of forming amyloid-like fibrils. Proc. Natl. Acad. Sci. U. S. A. 107, 3487–3492 (2010).

25. M. R. Sawaya, S. Sambashivan, R. Nelson, M. I. Ivanova, S. A. Sievers, M. I. Apostol, M. J. Thompson, M. Balbirnie, J. J. W. Wiltzius, H. T. McFarlane, A. Ø. Madsen, C. Riekel, D. Eisenberg, Atomic structures of amyloid cross-β spines reveal varied steric zippers. Nature. 447, 453–457 (2007).

26. S. A. Sievers, J. Karanicolas, H. W. Chang, A. Zhao, L. Jiang, O. Zirafi, J. T. Stevens, J. Münch, D. Baker, D. Eisenberg, Structure-based design of non-natural amino-acid inhibitors of amyloid fibril formation. Nature. 475, 96–100 (2011).

27. B. Neddenriep, A. Calciano, D. Conti, E. Sauve, M. Paterson, E. Bruno, D. A. Moffet, Short Peptides as Inhibitors of Amyloid Aggregation. Open Biotechnol. J. 5, 39–46 (2012).

28. A. Soragni, D. M. Janzen, L. M. Johnson, A. G. Lindgren, A. Thai-Quynh Nguyen, E. Tiourin, A. B. Soriaga, J. Lu, L. Jiang, K. F. Faull, M. Pellegrini, S. Memarzadeh, D. S. Eisenberg, A Designed Inhibitor of p53 Aggregation Rescues p53 Tumor Suppression in Ovarian Carcinomas. Cancer Cell. 29, 90–103 (2016).

29. P. M. Seidler, D. R. Boyer, J. A. Rodriguez, M. R. Sawaya, D. Cascio, K. Murray, T. Gonen, D. S. Eisenberg, Structure-based inhibitors of tau aggregation. Nat. Chem. 10, 170–176 (2018).

30. L. Saelices, K. Chung, J. H. Lee, W. Cohn, J. P. Whitelegge, M. D. Benson, D. S. Eisenberg, Amyloid seeding of transthyretin by ex vivo cardiac fibrils and its inhibition. Proc. Natl. Acad. Sci. U. S. A. 115, E6741–E6750 (2018).

31. S. L. Griner, P. Seidler, J. Bowler, K. A. Murray, T. P. Yang, S. Sahay, M. R. Sawaya, D. Cascio, J. A. Rodriguez, S. Philipp, J. Sosna, C. G. Glabe, T. Gonen, D. S. Eisenberg, Structure-based inhibitors of amyloid beta core suggest a common interface with tau. Elife. 8 (2019), doi:10.7554/eLife.46924.

32. S. Sangwan, S. Sahay, K. A. Murray, S. Morgan, E. L. Guenther, L. Jiang, C. K. Williams, H. V Vinters, M. Goedert, D. S. Eisenberg, Inhibition of synucleinopathic seeding by rationally designed inhibitors. Elife. 9 (2020), doi:10.7554/eLife.46775.

33. A. Leaver-Fay, M. Tyka, S. M. Lewis, O. F. Lange, J. Thompson, R. Jacak, K. Kaufman, P. D. Renfrew, C. A. Smith, W. Sheffler, I. W. Davis, S. Cooper, A. Treuille, D. J. Mandell, F. Richter, Y. E. A. Ban, S. J. Fleishman, J. E. Corn, D. E. Kim, S. Lyskov, M. Berrondo, S. Mentzer, Z. Popović, J. J. Havranek, J. Karanicolas, R. Das, J. Meiler, T. Kortemme, J. J. Gray, B. Kuhlman, D. Baker, P. Bradley, Rosetta3: An object-oriented software suite for the simulation and design of macromolecules. Methods Enzymol. 487, 545–574 (2011).

34. D. M. Fowler, A. V Koulov, C. Alory-Jost, M. S. Marks, W. E. Balch, J. W. Kelly, Functional Amyloid Formation within Mammalian Tissue. PLoS Biol. 4, e6 (2005).

35. D. M. Fowler, A. V Koulov, W. E. Balch, J. W. Kelly, Functional amyloid-from bacteria to humans. Trends Biochem. Sci. 32, 217–24 (2007).

36. C. L. L. Pham, A. H. Kwan, M. Sunde, Functional amyloid: widespread in Nature, diverse in purpose. Essays Biochem. 56, 207–219 (2014).

37. E. Tayeb-Fligelman, O. Tabachnikov, A. Moshe, O. Goldshmidt-Tran, M. R. Sawaya, N. Coquelle, J.-P. Colletier, M. Landau, The cytotoxic Staphylococcus aureus PSMα3 reveals a cross-α amyloid-like fibril. Science. 355 (2017).

38. N. Salinas, J. P. Colletier, A. Moshe, M. Landau, Extreme amyloid polymorphism in Staphylococcus aureus virulent PSMα peptides. Nat. Commun. 9, 3512 (2018).

39. N. Salinas, E. Tayeb-Fligelman, M. D. Sammito, D. Bloch, R. Jelinek, D. Noy, I. Usón, M. Landau, The amphibian antimicrobial peptide uperin 3.5 is a cross-α/cross-β chameleon functional amyloid. Proc. Natl. Acad. Sci. U. S. A. 118 (2021), doi:10.1073/pnas.2014442118.

40. D. T. Murray, M. Kato, Y. Lin, K. R. Thurber, I. Hung, S. L. McKnight, R. Tycko, Structure of FUS Protein Fibrils and Its Relevance to Self-Assembly and Phase Separation of Low-Complexity Domains. Cell. 171, 615–627.e16 (2017).

41. S. Brocca, R. Grandori, S. Longhi, V. Uversky, liquid–liquid phase separation by intrinsically disordered protein regions of viruses: Roles in viral life cycle and control of virus-host interactions. Int. J. Mol. Sci. 21, 1–31 (2020).

42. A. W. P. Fitzpatrick, B. Falcon, S. He, A. G. Murzin, G. Murshudov, H. J. Garringer, R. A. Crowther, B. Ghetti, M. Goedert, S. H. W. Scheres, Cryo-EM structures of Tau filaments from Alzheimer’s disease brain. Nature. 547, 185–190 (2017).

43. B. Li, P. Ge, K. A. Murray, P. Sheth, M. Zhang, G. Nair, M. R. Sawaya, W. S. Shin, D. R. Boyer, S. Ye, D. S. Eisenberg, Z. H. Zhou, L. Jiang, Cryo-EM of full-length α-synuclein reveals fibril polymorphs with a common structural kernel. Nat. Commun. 9, 3609 (2018).

44. P. M. Seidler, D. R. Boyer, K. A. Murray, T. P. Yang, M. Bentzel, M. R. Sawaya, G. Rosenberg, D. Cascio, C. K. Williams, K. L. Newell, B. Ghetti, M. A. DeTure, D. W. Dickson, H. V. Vinters, D. S. Eisenberg, Structure-based inhibitors halt prion-like seeding by Alzheimer’s disease-and tauopathy-derived brain tissue samples. J. Biol. Chem. 294, 16451–16464 (2019).

45. S. W. Roush, T. V. Murphy, M. M. Basket, J. K. Iskander, J. S. Moran, J. F. Seward, A. Wasley, Historical comparisons of morbidity and mortality for vaccine-preventable diseases in the United States. J. Am. Med. Assoc. 298, 2155–2163 (2007).

46. S. Lopez-Leon, T. Wegman-Ostrosky, C. Perelman, R. Sepulveda, P. A. Rebolledo, A. Cuapio, S. Villapol, medRxiv, in press, doi:10.1101/2021.01.27.21250617.

47. A. Lippi, R. Domingues, C. Setz, T. F. Outeiro, A. Krisko, SARS-CoV-2: At the Crossroad Between Aging and Neurodegeneration. Mov. Disord. 35, 716–720 (2020).

48. M. Goedert, D. S. Eisenberg, R. A. Crowther, Propagation of Tau Aggregates and Neurodegeneration. Annu. Rev. Neurosci. 40, 189–210 (2017).

49. J. C. Wootton, S. Federhen, Analysis of compositionally biased regions in sequence databases. Methods Enzymol. 266, 554–571 (1996).

50. P. M. Seidler, thesis, State University of New York at Buffalo (2014).

51. K. L. Heckman, L. R. Pease, Gene splicing and mutagenesis by PCR-driven overlap extension. Nat. Protoc. 2, 924–932 (2007).

52. M. Faller, M. Matsunaga, S. Yin, J. A. Loo, F. Guo, Heme is involved in microRNA processing. Nat. Struct. Mol. Biol. 14, 23–29 (2007).

53. W. Kabsch, XDS. Acta Crystallogr. Sect. D Biol. Crystallogr. 66, 125–132 (2010).

54. A. J. McCoy, R. W. Grosse-Kunstleve, P. D. Adams, M. D. Winn, L. C. Storoni, R. J. Read, IUCr, *Phaser* crystallographic software. J. Appl. Crystallogr. 40, 658–674 (2007).

55. G. M. Sheldrick, Experimental phasing with SHELXC/D/E: Combining chain tracing with density modification. Acta Crystallogr. Sect. D Biol. Crystallogr. 66, 479–485 (2010).

56. G. N. Murshudov, P. Skubák, A. A. Lebedev, N. S. Pannu, R. A. Steiner, R. A. Nicholls, M. D. Winn, F. Long, A. A. Vagin, REFMAC5 for the refinement of macromolecular crystal structures. Acta Crystallogr. Sect. D Biol. Crystallogr. 67, 355–367 (2011).

57. P. Emsley, B. Lohkamp, W. G. Scott, K. Cowtan, Features and development of Coot. Acta Crystallogr. Sect. D Biol. Crystallogr. 66, 486–501 (2010).

58. The PyMOL Molecular Graphics System, Version 1.2r3pre, Schrödinger, LLC.

59. E. B. Saff, A. B. J. Kuijlaars, Distributing many points on a sphere. Math. Intell. 19, 5–11 (1997).

60. M. C. Lawrence, P. M. Colman, Shape complementarity at protein/protein interfaces. J. Mol. Biol. 234, 946–950 (1993).

61. M. D. Winn, C. C. Ballard, K. D. Cowtan, E. J. Dodson, P. Emsley, P. R. Evans, R. M. Keegan, E. B. Krissinel, A. G. W. Leslie, A. McCoy, S. J. McNicholas, G. N. Murshudov, N. S. Pannu, E. A. Potterton, H. R. Powell, R. J. Read, A. Vagin, K. S. Wilson, Overview of the *CCP* 4 suite and current developments. Acta Crystallogr. Sect. D Biol. Crystallogr. 67, 235–242 (2011).

